# The Spectre of Too Many Species

**DOI:** 10.1101/263996

**Authors:** Adam D. Leaché, Tianqi Zhu, Bruce Rannala, Ziheng Yang

**Author notes:** Correspondence to be sent to: Ziheng Yang, Department of Genetics, University College London, London WC1E 6BT, UK. Those authors contributed equally to the study (Adam D. Leaché, Tianqi Zhu).

## Abstract

Recent simulation studies examining the performance of Bayesian species delimitation as implemented in the BPP program have suggested that BPP may detect population splits but not species divergences and that it tends to over-split when data of many loci are analyzed. Here we confirm several of these results and provide their mathematical justifications. We point out that the distinction between population and species splits made in the protracted speciation model has no influence on the generation of gene trees and sequence data, which explains why no method can use such data to distinguish between population splits and speciation. We suggest that the the protracted speciation model is unrealistic and its mechanism for assigning species status contradicts prevailing taxonomic practice. We confirm the suggestion, based on simulation, that in the case of speciation with gene flow, Bayesian model selection as implemented in BPP tends to detect population splits when the amount of data (the number of loci) increases so over-splitting is a legitimate concern. We discuss the use of a recently proposed empirical genealogical divergence index (*gdi*) for species delimitation and illustrate that parameter estimates produced by a full likelihood analysis as implemented in BPP provide much more reliable inference under the *gdi* than the approximate method PHRAPL. We suggest that the Bayesian model-selection approach is useful for identifying sympatric cryptic species while Bayesian parameter estimation under the multispecies coalescent can be used to implement empirical criteria for determining species status among allopatric populations.

---

In the past decade, the multispecies coalescent (MSC) model (Rannala and Yang, 2003) has emerged as an important framework for statistical analysis of genomic sequence data from closely related species. Under the model, different genomic regions (called loci) may have different genealogical histories due to coalescent processes occurring in the extinct ancestral species. The MSC thus naturally accommnodates gene tree heterogeneity across the genome. Likelihood-based inference under the MSC averages over the gene trees for multiple loci, achieved through either numerical integration (Yang, 2002; Zhu and Yang, 2012) or Bayesian Markov chain Monte Carlo (MCMC) (Edwards, 2009; Heled and Drummond, 2010; Yang and Rannala, 2010, 2014). Averaging over gene trees incurs a heavy computational burden but has the benefit of accommodating phylogenetic uncertainty at individual loci, which is important when the species are closely related and the sequence alignment at each locus has low phylogenetic information content (Xu and Yang, 2016). Given the species phylogeny, the MSC can be used to estimate important parameters concerning species divergences, such as the population sizes of modern and ancestral species, species divergence times, and past migration patterns and rates (Burgess and Yang, 2008; Hey, 2010; Mailund *et al.*, 2012; Takahata *et al.*, 1995). The MSC also provides the appropriate inference framework for estimating species phylogenies while accommodating gene tree heterogeneity caused by deep coalescence and incomplete lineage sorting (Edwards, 2009; Heled and Drummond, 2010; Maddison, 1997; Nichols, 2001; Yang and Rannala, 2014). It has been applied to species identification (assignment) and found to achieve better statistical performance than DNA barcoding based on a simple distance threshold (Yang and Rannala, 2017). The MSC has also been used to address the problem of species discovery (or delimitation) (Yang and Rannala, 2010, 2014). Different species delimitation models are formulated as competing statistical models and inferred from the genetic data through Bayesian model selection (i.e., through calculation of posterior model probabilities). Species delimitation is a complex issue, however, partly because there is no universally accepted definition of species (Mallet, 2013).

Two recent studies (Jackson *et al.*, 2017; Sukumaran and Knowles, 2017) used computer simulation to evaluate the performance of Bayesian species delimitation as implemented in the software package BPP (Bayesian Phylogenetics and Phylogeography) (Rannala and Yang, 2013; Yang and Rannala, 2010). Both studies concluded that BPP may over-split, capturing population splits rather than species divergences. Sukumaran and Knowles (2017) simulated phylogenies, gene trees and sequence data under the protracted speciation model (PSM) (Etienne *et al.*, 2014; Rosenblum *et al.*, 2012) which distinguishes between populations (incipient species) and species. They concluded that in some cases BPP delimited population structure rather than species. Jackson *et al.* (2017) simulated sequence data under the MSC model on a given species tree, and then used a heuristic genealogical divergence index (*gdi*) to define species status. They found that their simulation-based heuristic method PHRAPL was more successful in inferring species status than BPP, which tended to split subdivided populations into species even in the face of high gene flow.

Here we examine the conditions of the simulations of Sukumaran and Knowles (2017) and Jackson *et al.* (2017) to evaluate the performance of BPP. Two features of the simulation of Sukumaran and Knowles (2017) are noteworthy. First, the species conversion process overlies the population branching process and is Markovian (memoryless) so the rate of species conversion (from incipient species to species) is fixed and independent of the duration of genetic isolation between incipient species. Moreover, the PSM distinguishes between populations and species but the species status of lineages is ignored when the gene trees and sequence data are generated under the MSC model for subsequent analysis using BPP. Second, the assignment of species status in the PSM does not appear to be consistent with current taxonomic practices or with most models of speciation.

In Jackson *et al.* (2017), a heuristic criterion was used to define species and that definition was used in PHRAPL but not in BPP when both programs were used to infer species status. We perform a fair comparison in which the same heuristic species definition is used in both PHRAPL and BPP analyses. We demonstrate that even though BPP ignores gene flow and is based on the simplistic JC mutation model (Jukes and Cantor, 1969), it provides more accurate parameter estimates and inference of species status than PHRAPL when both programs use the same heuristic definition of species. The large sample properties of Bayesian species delimitation are described using new asymptotic results for the statistical behavior of BPP in delimiting species as the number of loci increases.

## PROTRACTED SPECIATION?

A defining feature of the simulation by Sukumaran and Knowles (2017) under the PSM is that the events that transform populations into species are independent of the process of genetic divergence among populations and the generation of gene trees and sequence data. The PSM distinguishes between populations and species but when the population tree is used to simulate gene trees and sequences no such distinction is made. The simulation could be viewed as representing the use of the neutral genome or noncoding DNA to delineate species boundaries.

### The likelihood principle

Sukumaran and Knowles (2017) observed that the BPP program cannot distinguish between populations and true species when species conversion rates are low. This inability of the BPP program (or more precisely of the sequence data) to distinguish between incipient and true species is expected and obvious without simulations. It is a direct consequence of the likelihood principle in statistics. This states that all information about the competing models and model parameters is contained in the likelihood function, which is the probability of the data given the model and parameters (O’Hagan and Forster, 2004, pp. 61-64). If two models make the same probabilistic predictions about the observable data and thus have identical likelihoods for all possible data outcomes, the models are not identifiable and the data cannot be used to distinguish them. Consider a coin-tossing experiment, in which a sophisticated probabilistic model is used to decide whether one should pray before every toss. As praying does not affect the outcome of the coin toss, the data of heads and tails will not contain any information about whether each coin toss is magical (preceded by prayer) or ordinary (without prayer). In a Godless universe the magic- and ordinary-toss models make exactly the same predictions about heads and tails, which consequently cannot be used to distinguish those models. The species conversion process is analogous to the prayer; it does not alter the likelihood of the observed genetic data and is therefore not identifiable.

### Going to extremes

The Sukumaran and Knowles (2017) model is extreme in several respects and is unlikely to be realistic for the majority of speciation processes in nature. The model posits an exaggerated form of punctuated equilibrium – exponentially distributed periods of stasis followed by an instantaneous conversion to a new species. At the conversion event, the new population and the parental population (which is only one generation older) are deemed distinct species; few species appear to have originated in this way. An alternative “gradualist” model would treat the morphological characters involved in species classification as quantitative traits that evolve according to a diffusion model determined by the effects of underlying mutational changes and genetic drift of allele frequencies. Two populations are recognized as different species if the difference in mean trait values exceeds some threshold. The mean difference that is chosen reflects the biologists perception of what species are and how morphologically distinct they should be. Under such a model there will be a strong covariance between genetic isolation, population divergence time and species status so that delimitation methods such as BPP should perform better. The gradualist model is, of course, another extreme and a more realistic model would include both morphological “jumps” and “diffusions” (see Discussion).

The way in which the PSM assigns species status is also problematic, contradicting prevailing taxonomic practices. In Figure 1 of Sukumaran and Knowles (2017), the different colors on branches signify distinct species produced by conversion events under the PSM (Fig. 1). It is possible for the model to generate species near the tips of the species tree, say, < 10 generations ago. However, taxonomists would not recognize recent divergences of only a few generations as valid speciation events. Instead, speciation is a consequence of an extended process of genetic isolation, and species status is assigned retrospectively based on empirical measures of morphological and/or genetic divergence. It is not possible to simulate species forward in time because the criterion of the systematist depends on the level of divergence between populations and this is only known after the simulation is completed (assuming that species do not arise instantaneously). To conclude, the PSM is unrealistic in several respects limiting its utility for evaluating species delimitation methods. In particular, its application may not justify the authors’ conclusion that MSC methods “delimit structure not species.”

**FIGURE 1.**
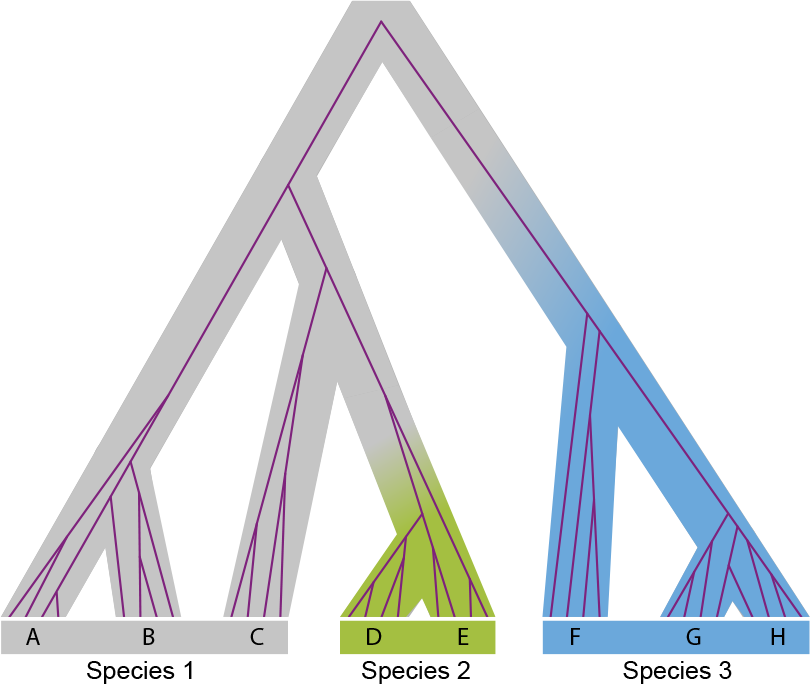
Figure 1 of Sukumaran and Knowles (2017) redrawn to illustrate the simulation of species (indicated by tip labels) under the protracted speciation model. The species tree is shown with one embedded gene tree (purple); protracted speciation events happen when the species tree changes color.

### Is a new delimitation method needed?

The PSM does not require the development of new approaches for delimitation. Implementing the PSM in BPP requires only a modification to the prior on species delimitation models used by the program. The likelihood of the sequence data under the MSC remains unchanged. Let *π_i_* be the prior for species delimitation model *i* and *M_i_* its marginal likelihood, which is the probability of the sequence alignments at multiple loci, averaged over the gene trees and over the parameters of the MSC model (*τs* and *θs*) through the prior. The posterior probability for delimitation model *i* is then

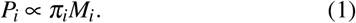

If we change the prior to 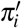, the new posterior will be

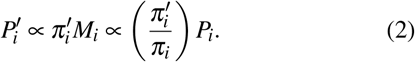

The use of the new prior simply modifies the posterior by a correction factor 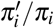, although the posteriors for all models need to be rescaled to sum to 1. Thus, the PSM can be specified by using a new prior (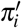) just like other priors on species delimitation models already implemented (Yang, 2015). From this argument, it is clear that the posterior probabilities for the species delimitation models can be extremely sensitive to the prior. It is equally obvious that the PSM, even if correctly implemented, will not be useful for reliable delimitation of species status, in the same sense that a correctly implemented model of stochastic prayers (e.g., a binomial model) will not allow us to use counts of heads and tails to distinguish between magic and ordinary coin tosses with any reliability.

## ASYMPTOTIC BAYESIAN SPECIES DELIMITATION

Jackson *et al.* (2017) simulated data under the MSC model with migration (Hey, 2010, the so-called isolation-with-migration or IM model) for two species/populations and analyzed them using BPP to calculate the posterior probabilities for the one-species and two-species models. They observed that the posterior probability for the two-species model increases when the number of loci increases. Here we investigate the asymptotic behavior of Bayesian posterior model probabilities and confirm that this is the expected behavior of Bayesian model selection and of the program.

### Choosing among wrong models

The asymptotic dynamics of Bayesian model selection depends on how wrong the two competing models are relative to the true data-generating model (Yang and Zhu, 2018). Here we consider independent and identically distributed (i.i.d.) models only, under which the data points *x_i_* (*i* = 1,…*L*) are i.i.d., with *x_i_* ~ *q*(*x_i_*). Let *X* = {*x_i_*}. The distance from any model *p*(*x*|*ϕ*) with parameters *ϕ* to the true model *q* is measured by the Kullback-Leibler (K-L) divergence

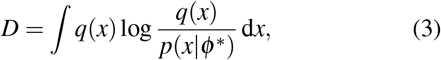

where *ϕ^*^* is the limiting maximum likelihood estimate (MLE) of *ϕ* under the model when the data size *L* → ∞, and is known as the best-fitting parameter value under the model (White, 1982). The K-L divergence *D* = 0 if the model encompasses the true model (or, in other words, is true), and *D* > 0 if the model is wrong.

Here the true model *q* is the MSC model with migration (the IM model). Under the model, the gene trees and sequence alignments are i.i.d. among loci, so that the datasize is the number of loci (*L*). Currently BPP does not accommodate migration or introgression and implements the complete isolation model only. The two models under comparison are then the one-species model (*H*_1_) with a single population-size parameter *ϕ*_1_ = {θ} and the two-species model (*H*_2_) with parameters ϕ_2_ = {τ, *θ_A_*, *θ_B_*, *θ_AB_*}, where *τ* (for *τ_AB_*) is the divergence time between the two species, and the *θ* s are the population size parameters for the two modern species *A* and *B* and for the ancestral species *AB,* with *θ* = 4*Nμ* (Fig. 2a). Both θ and τ are measured in the expected number of mutations per site. As the true model involves migration, both models *H*_1_ and *H*_2_ are wrong, with *D*_1_ > 0, *D*_2_ > 0. Note that *H_1_* is a special case of *H*_2_ since the two models are equivalent when *τ* = 0 in *H*_2_, in which case parameters *θ_A_* and *θ_B_* in *H_2_* are unidentifiable. The dynamics of the posterior probabilities for *H*_1_ and *H*_2_ depends on whether *H*_1_ and *H*_2_ are equally wrong (in which case *D*_1_ = *D*_2_ > 0) or *H*_1_ is less wrong than *H*_1_ (with *D*_1_ > *D*_2_ > 0), or equivalently on whether the best fitting parameter value for *τ* in *H*_2_ is

**FIGURE 2.**
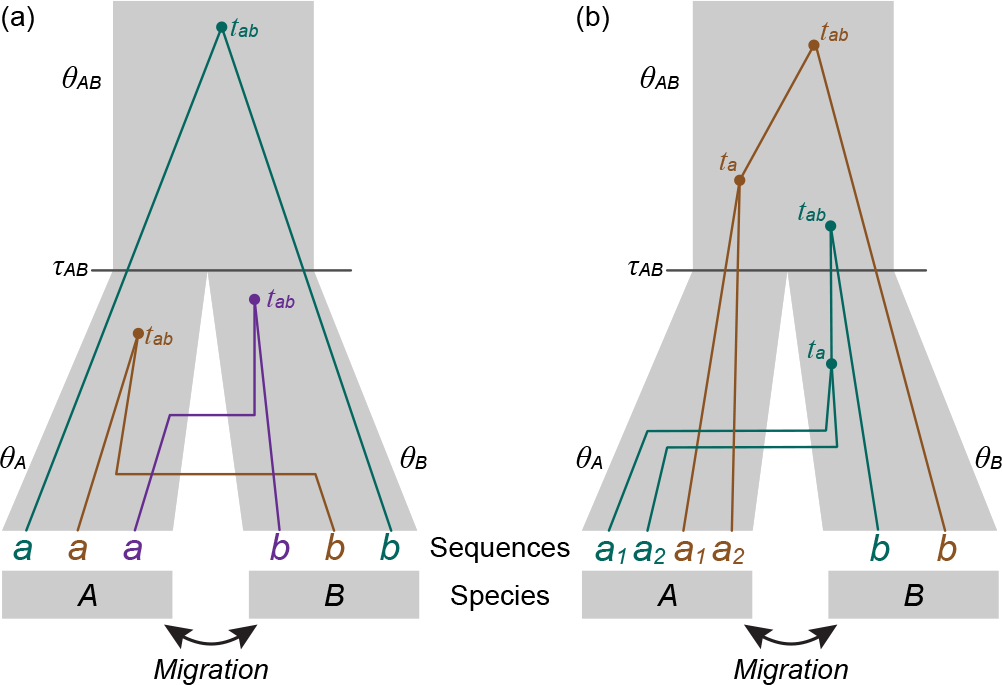
(a) A species tree for two species (*A* and *B*) and three gene trees for two sequences (*a* and *b*), used to illustrate the asymptotics of Bayesian model selection. The coalescence between the two sequences occurs before species divergence in the brown and purple gene trees (with *t* < τ) and after in the green gene tree (with *t* > τ). (b) A species tree for two species (*A* and *B*) and two gene trees for three sequences (*a*_1_ and *a*_2_ from species *A* and *b* from species *B*), used to illustrate the computation of the *gdi.* Both gene trees have the same topology *G*_1_ = ((*a*_1_, *a*_2_), but the coalescence between *a*_1_ and *a*_2_ occurs before species divergence (in species *B*) in the green tree (with *t_a_* < *τ_AB_*) and after in the brown tree (with *t_a_* > *τ_AB_*).

*τ^*^* = 0 or > 0. If *τ*^*^ = 0, the two models will be equally wrong, and they are also unidentifiable in the limit of infinite data. Then *H*_1_, with fewer parameters, dominates, with its posterior probability approaching 100% when the number of loci *L* increases. In contrast, if *τ^*^* > 0, *H*_2_ is less wrong than *H*_1_, and *H*_2_ will dominate. While an analytical proof is not available, we analyze increasingly larger datasets to examine the asymptotic behavior of the MLEs numerically. Our calculations suggest that the second case applies: when the true model is the MSC model for two populations with migration, the two-species isolation model is less wrong than the one-species model and dominates in the posterior when the number of loci increases.

As an example, we simulate large datasets with many loci, each of 500 sites, under the symmetrical IM model for two species with τ = 0.01 for the species divergence and *θ_A_ = θ_B_ = θ_AB_ = θ* = 0.01 for all populations, and with migration rates between the two populations to be *M_AB_ = M_BA_ = M = N_m_* = 10 immigrants per generation (Fig. 2a). In this paper, the (scaled) migration rate is defined as *M_i,j_ = N_j_m_ij_*, the expected number of immigrants in population *j* from population *i* per generation, with *m_ij_* to be the proportion of immigrants in population *j.* The MCCOAL program, in the BPP package, was used to generate gene trees and sequence alignments under the JC model (Jukes and Cantor, 1969). Each locus has two sequences, *a* and *b*, from species *A* and *B*, respectively. At those parameter values, sequences *a* and *b* coalesce before species divergence (with *t < τ*, as in the brown and purple gene trees of Fig. 2a) at 62.75% of loci, which is very similar to the probability for *t* < τ (63.21%) if the two sequences are from the same population.

The data are then analyzed using the 3S program to obtain the MLEs for the two parameters (*θ_AB_* and *τ*) under the two-species MSC model with no migration (*H*_2_) (Dalquen *et al.*, 2017; Yang, 2002). The estimate of *θ_AB_* is 0.0158. The MLE t ranged from 0.00033-0.00036 over ten replicates for *L* = 10^5^ and over 0.000329-0.000348 for *L =* 2 × 10^5^. Based on the stability of the estimates among the replicate datasets and between the large values of L, we suggest that at the limit of infinitely many loci, the best-fitting parameter value is *τ^*^* = 0.00034. We note that the best-fitting parameter value depends on the configuration of the data such as the number of sequences per locus and the number of sites, as well as the parameters of the MSC model with migration (*τ*s, *θ*s, and *M*’s). If the sequence length is 250 sites instead of 500, we obtain *τ*^*^ ~ 0.00062 instead of 0.00034. Those results provide numerical evidence that at the limit of infinite data, *τ*^*^ > 0, so that the two-species model will dominate the posterior, even though the migration rates are so high between the two populations that they should be considered one species by any species definition.

### Including a migration model

Note that if Bayesian model selection is conducted under the IM model, incorporating migration, the two-species model with migration will be correct (with *D*_2_ = 0), while the one-species model will be wrong (with *D*_1_ > 0). Then the two-species model will dominate with the posterior probability approaching 100% as the number of loci increases. This is the case even if the migration rate *M = Nm* is very large (but finite). Thus if we use Bayesian model selection to infer species status (treating a population split as a speciation event) then incorporating migration into the MSC model will not correct the problem of over-splitting.

In conclusion, the concern that Bayesian model selection as implemented in BPP may over-split and recognize too many species in subdivided populations with ongoing gene flow is legitimate. Over-splitting may be of particular concern when hundreds or thousands of loci are analyzed. If two populations are truly panmictic, the model with fewer parameters will be favored, and the populations will be correctly lumped into one species. However, if there is partial subdivision (even with relatively high levels of gene flow) the method will prefer the two species model asymptotically as the number of loci increases. One possible solution is to include a model with gene flow and use model selection to choose among 3 models: (1) a single population; (2) two completely isolated populations; and (3) two populations with gene flow. A choice of model 1 strongly suggests a single species; a choice of model 2 suggests two species but a final decision should be based on a consideration of the population divergence time and other relevant information (morphology, etc); a choice of model 3 allows either one species or 2, depending on considerations such as the degree of gene flow, distinctness of morphology, and so on.

## HEURISTIC SPECIES DELIMITATION

Jackson *et al.* (2017) suggested a heuristic criterion for species delimitation based on a genealogical divergence index (*gdi*) between populations that can be calculated using estimates of parameters under the MSC model with migration (*τ*, *θ*, and *M*). Suppose one samples two sequences (*a*_1_ and *a*_2_) from population *A* and one sequence (*b*) from population *B* (see Fig. 2b). Let the probability that the two sequences from population *A* coalesce first, so that the gene tree is *G*_1_ = ((*a*_1_,*a*_2_),*b*), be

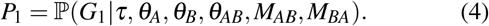

Obviously *P*_1_ ranges from 1/3 (when the three sequences are interchangeable, as in the case of *M_AB_ = M_AB_ = ∞*) to 1. Jackson *et al.* (2017) rescaled *P*_1_ so that the genealogical divergence index,

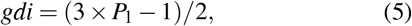

ranges from 0 to 1 when *P*_1_ goes from 1/3 to 1. In the special case of no migration (with *M_AB_ = M_BA_* = 0), we have *P*_1_ = 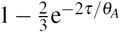 and

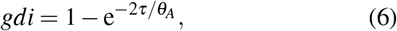

where 2*τ/θ_A_* is the population divergence time in coalescent units (with one coalescent time unit to be 2*N_A_* generations) and 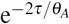 is the probability that the two sequences from population *A* (*a*_1_ and *a*_2_) do not coalesce before reaching species divergence (τ) when we trace the genealogy backwards in time.

### The gdi heuristic for species identification

Jackson *et al.* (2017) calculated the *gdi* as defined in equations 4 and 5 by simulating 10,000 gene trees under the MSC model with migration. Here we provide its analytical computation, using the Markov chain characterization of the coalescent process with migration (Dalquen *et al.* 2017; Hobolth *et al.* 2011; Zhu and Yang 2012). For two populations (*A* and *B*) with gene flow and three sequences (*a*_1_, *a*_2_, and *b*), the genealogical process of coalescent and migration when one traces the history of the sample backwards in time can be described by a Markov chain with 21 states. The state of the chain is specified by the number of sequences remaining in the sample and the populations in which they reside, or by the population IDs (*A* and *B*) and the sequence IDs (*a*_1_, *a*_2_), *b* etc.). For example, the state *A_a1_A_a2_B_b_* means that the three sequences *a*_1_, *a*_2_ and *b* are in populations *A, A*, and *B*, respectively. We also write this as ‘*AAB*’. This is the initial state. State *A*_*a*_1_*a*_2__*B*_*b*_, abbreviated ‘*AB_b_*, means that two sequences remain in the sample, with the ancestor of sequences *a*_1_ and *a*_2_ in population *A* and sequence *b* in population *B.*

The transition rate matrix of the Markov chain *Q* = {*q_ij_*} is given in table 1. The transition probability matrix over time *t* is then *P*(*t*) = {*p_ij_*(*t*)} = *e*^*Qt*^, where *p_ij_*(*t*) is the probability that the Markov chain is in state *j* at time *t* in the past given that it is in state *i* at time 0 (the present time). Suppose *Q* has the spectral decomposition

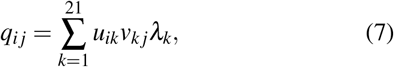

where *λ_k_* are the eigenvalues of *Q*, columns in *U = {u_ij_}* are the corresponding right eigenvectors, and rows in *V* = {*v_i,j_* = *U*^-1^ are the left eigenvectors. Then

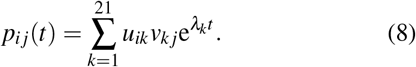

Gene tree *G*_1_ = ((*a*_1_, *a*_2_), *b*) can be generated in two ways. The first is for sequences *a*_1_ and *a*_2_ to coalesce before reaching the ancestral population, with *t* < *τ* (as in the green gene tree of Fig. 2b). Sequence *b* then joins the ancestor of sequences *a*_1_ and *a*_2_ either before species divergence at *τ*, in which case the root of the gene tree is younger than species divergence, or after, in which case the root of the gene tree is older than *τ* (the latter case is illustrated in the green gene tree of Fig. 2b).

The probability density that sequences *a*_1_ and *a*_2_ coalesce at time *t* < *τ* is given by

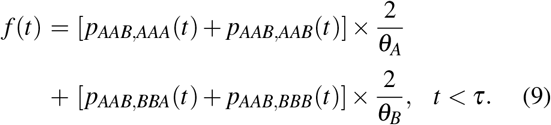

This is a sum of two terms, corresponding to the first coalescent (between sequences *a*_1_ and *a*_2_) occurring in populations *A* and *B*, respectively. The first term is the probability, *p_AAB,AAA_*(*t*) + *p_AAB,AAB_*(*t*), that sequences *a*_1_ and *a*_2_ are in population *A* right before time *t*, times the rate for them to coalesce (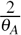). Similarly the second term is the probability density that sequences *a*_1_ and *a*_2_ coalesce at time *t* in population *B* (Fig. 2b, green gene tree).

The second way of generating gene tree *G*_1_ is for sequences *a*_1_ and *a*_2_ to coalesce after population divergence, with *t* > *τ* (as in the brown gene tree of Fig. 2b). This occurs with probability 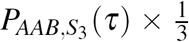, where *S*_3_ = {*AAA,AAB,ABA,ABB,BAA,BAB,BBA,BBB*} is the set of states with three sequences, and 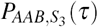 is the probability that no coalescent event occurs during the time interval (0, *ϕ*). In this scenario, the gene tree root must be older than *ϕ*.

Thus combining the two possibilities for generating gene tree *G*_1_, we have

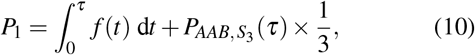

where *f*(*t*) is given in equation 9. To calculate the integral in equation 10, note that from equation 8,

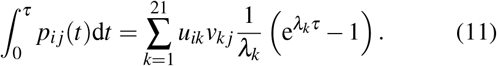

We calculated *P*_1_ under the symmetrical migration model with *θ_A_ = θ_B_ = θ* and *M_AB_* = *M_BA_ = M = Nm.* Figure 3b shows P_1_ plotted against 2*τ/θ* (population divergence time in coalescent units) and *M* under the symmetrical migration model. This is a more accurate calculation than Fig. 6 of Jackson *et al.* (2017), which was based on simulating gene trees, even though the two approaches are equivalent if a huge number of replicates is used in the simulation.

**FIGURE 3.**
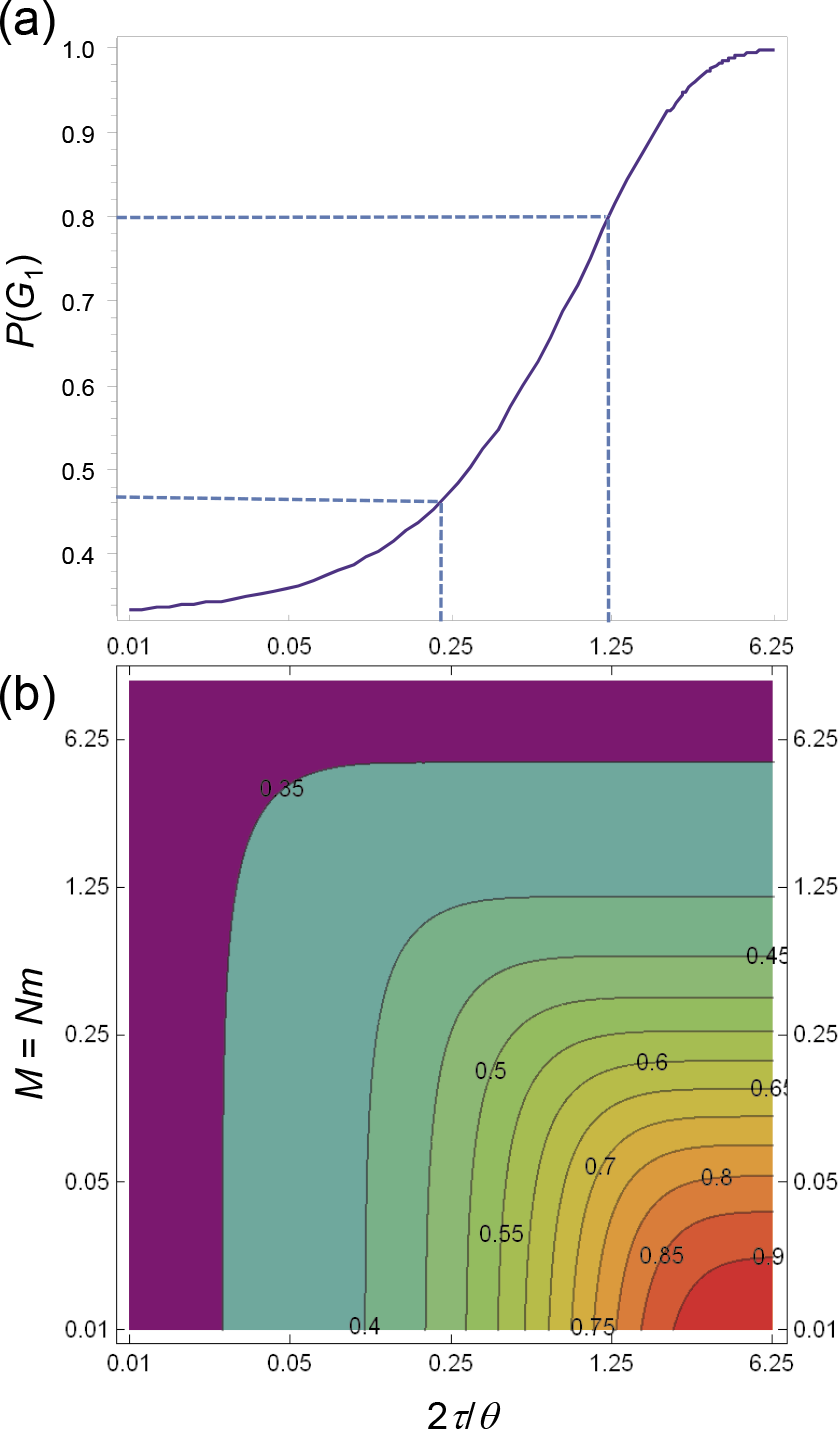
Probability 핡(*G*_1_) of gene tree *G*_1_ = ((a_1_,*a*_2_),*b*), plotted (a) as a function of population divergence in coalescent units (2*τ/θ*) in a pure isolation model for two populations without gene flow, and (b) as a function of population divergence in coalescent units (2*τ/θ*) and scaled migration rate *M = Nm.* According to Jackson *et al.* (2017), the lower and upper limits of *P*_1_ for species delimitation are 0.47 and 0.8.

Based on the meta-analysis of Pinho and Hey (2010), Jackson *et al.* (2017) suggested the rule of thumb that *gdi* values < 0.2 suggest a single species and *gdi* values > 0.7 suggest distinct species, while *gdi* values within the range indicate ambiguous delimitation. The limits of 0.2 and 0.7 for *gdi* correspond to 0.47 and 0.8 for *P*_1_, and, in the case of no migration, to 0.22 and 1.20 for the population divergence in coalescent units (2*τ/θ*) (Fig. 3a).

## SUBJECTIVELY DEFINED SPECIES

Jackson *et al.* (2017) simulated data under the MSC model with migration for two populations and analyzed the data using PHRAPL and BPP. While the true model used in the simulation always had two populations, the *gdi* was used to define species status. This criterion was used in the PHRAPL analysis of the simulated data to infer species status, but not in BPP. It was then found that PHRAPL out-performed BPP (Jackson *et al.* 2017, Fig. 4), and that BPP tended to over-split, identifying too many species.

### A fair comparison

Both BPP and PHRAPL can estimate the parameters of the MSC model, although PHRAPL accommodates gene flow while BPP in its current implementation assumes no gene flow. Here we apply the *gdi* definition of species status in BPP, so that the same criterion is used by BPP and PHRAPL. A simple approach is to use the posterior means of the parameters under the MSC generated by BPP to calculate the *gdi* (equation 6). We use this method here. A more sophisticated approach, which we use later in the analysis of the empirical datasets, is to generate a posterior distribution of *gdi* using the sample of parameters taken during the MCMC.

We thus repeated the simulation of Jackson *et al.* (2017, fig. 4), applying *gdi* to BPP parameter estimates. The true species tree is (*(A,B),C*), with six sets of species divergence time parameters, with *τ_AB_* = 0.05*θ*,0.125*θ*,0.25*θ*,1*θ*,2*θ*, 4*θ*, and *τ_ABC_* = 2.5*θ*, 2.5*θ*,2.5*θ*,2.5*θ*,5*θ*,10*θ*, with *θ =* 0.005. Note that *τ_ABC_* is much larger than *τ_AB_*, so that species *C* is a distant outgroup, and the focus is on whether populations *A* and *B* are one or two species. Migration is assumed to occur between *A* and *B*, with 4*Nm* = 0,0.5,2, and 5, where *Nm* is the number of immigrants per generation. The sequence data were simulated under the HKY model (Hasegawa *et al.*, 1985), with base frequencies 0.3, 0.2, 0.3, and 0.2 (for T, C, A, and G) and transition/transversion rate ratio *κ* = 3. For each of the 4 × 6 parameter combinations for *M* and *τ*, 50 replicate datasets were simulated. There are 50 loci in each dataset, with 20 sequences from each of the three species, and 500 sites in the sequence. The data were simulated using the MCCOAL program, part of the BPP release, as detailed in Zhang *et al.* (2011). We used BPP version 4.0 to estimate the parameters in the MSC model on the fixed species tree ((*A,B),C*) (this is the A00 analysis of Yang, 2015). Version 4.0 of the program assigns inverse-gamma priors on parameters. We used the shape parameter 3 in the inverse-gamma priors, while the prior means are set to match the true values: *θ* ~ IG(3, 0.01) with mean 0.01/(3 – 1) = 0.005, and *τ_ABC_* ~ IG(3, 0.025), IG(3, 0.05), and IG(3, 0.1), for the three true *τ_ABC_* values. Note that the value 3 for the shape parameter means that the inverse-gamma priors are diffuse, with the coefficient of variation to be 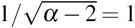. Estimation of parameters under the MSC is known to be fairly robust to the priors, for example, to a one order-of-magnitude change to the prior means (Burgess and Yang, 2008). After BPP generated the posterior distribution of the parameters, we used the posterior means to calculate *gdi* using equation 6, with *τ* = *τ_AB_* and *θ* = (*θ_A_* + *θ_B_*)/2.

The results are shown in figure 4. Even though it ignores migration and uses an overly simplistic JC mutation model (while the true model is HKY), BPP performed better than PHRAPL in delimiting species status defined by the *gdi*, especially at high migration rates (with 4*Nm =* 2 or 5). This result may seem counterintuitive, since the data were simulated with migration and PHRAPL allows for migration so that there is no model violation, while BPP ignores migration so that its model is violated.

### Shortcomings of approximate methods

We suggest that two factors may account for the poorer performance of PHRAPL in this simulation. First, PHRAPL is a summary method for estimating parameters, and it relies on gene tree topologies and ignores branch lengths. As a result, parameter estimates may be biased or even inconsistent due to phylogenetic errors of gene tree reconstruction (Yang, 2002). Second, use of the gene tree topologies while ignoring the branch lengths leads to information loss and may even cause identifiability problems. In the simple case of three species and three sequences, with one sequence from each species, there is only one degree of freedom in the data of gene tree topologies, which is the proportion of the most common gene tree topology. Under the complete-isolation model (with *M =* 0), this allows one to estimate the internal branch length on the species tree in coalescent units, 2(*τ_ABC_* – *τ_AB_)/θ_AB_*, while other parameters in the model are unidentifiable (Xu and Yang, 2016). Even the internal branch length is estimated inconsistently because phylogenetic reconstruction errors tend to inflate gene tree-species tree mismatches (Yang, 2002).

**FIGURE 4.**
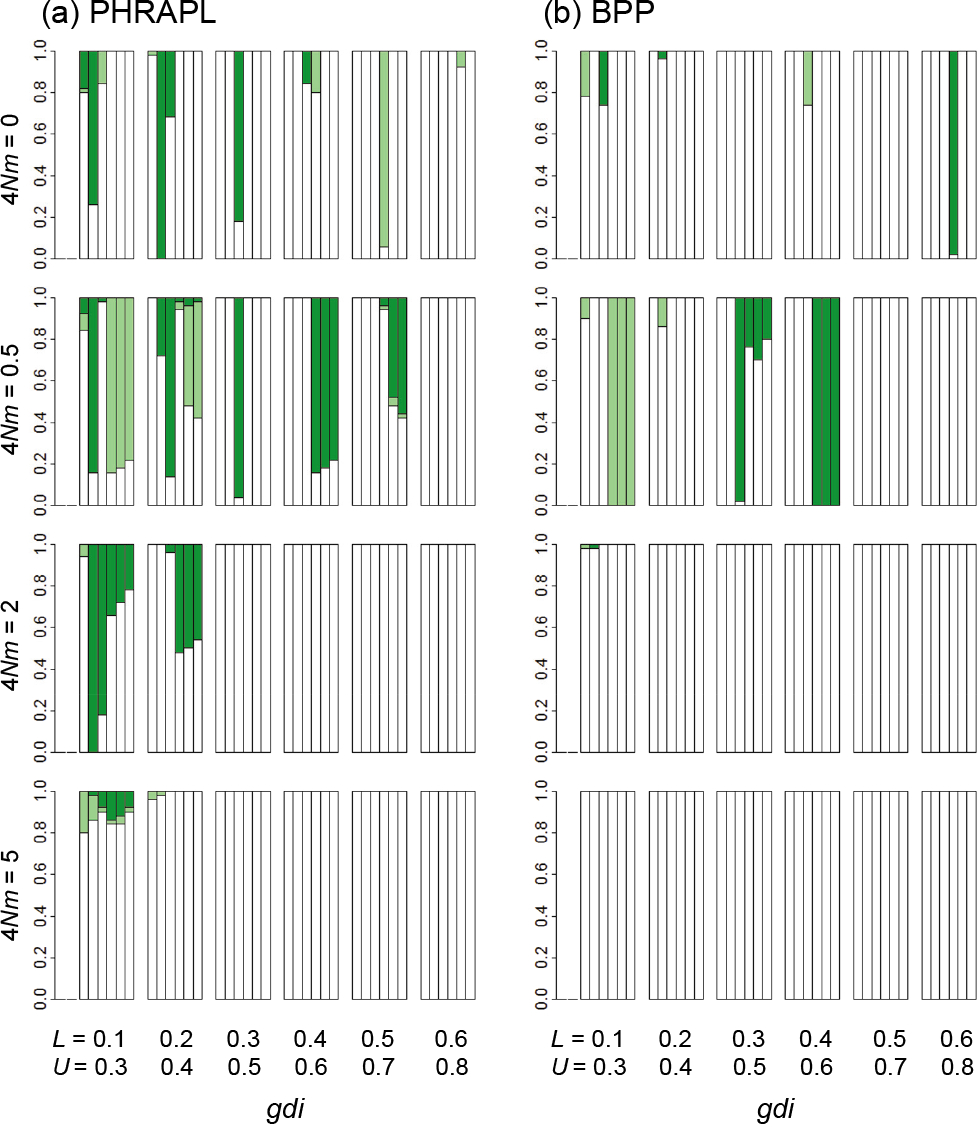
Accuracy of species delimitation using the *gdi* with parameters estimated from data of 50 loci using (a) PHRAPL and (b) BPP. Species status is defined using the *gdi* at different cutoffs (*L* and *U).* This is calculated by simulating 10,000 gene trees under the MSC model with migration for PHRAPL, and analytically for BPP. Along the x-axis, each group of bars gives results for different *gdi* cutoffs. Below the lower bound (L), populations *A* and *B* are defined as a single species; above the upper bound (*U), A* and *B* are defined as separate species, while between the bounds, the species status is ambiguous. The six bars within each group represent the six sets of species divergence times (*ϕ*s). The bar shadings are white = the inferred delimitation outcome matched the true outcome; light green/gray = ambiguity was inferred when the true delimitation is known (insufficient power); dark green/gray = delimitation was inferred (whether one or two species) when the truth was ambiguous (excessive confidence); and black = one species was inferred when there were two, or vice versa. The results for PHRAPL are recreated using the R code from Jackson *et al.* (2017, Fig. 4).

The cases with more than three sequences per locus and with migration may be more complex, but it should not be surprising that approximate methods that rely on summary statistics such as gene tree topologies will suffer from an information loss. In contrast, BPP is a full-likelihood method and makes use of information in the gene tree branch lengths (coalescent times) as well as topologies, while accommodating phylogenetic uncertainties due to the limited number of informative sites at each locus (Xu and Yang, 2016; Yang, 2014). Even though BPP operates under a wrong model that ignores migration, the sequence data at multiple loci may be informative about the expected gene tree configurations. Nevertheless, extension of BPP to allow for gene flow will provide more accurate estimation of parameters in the MSC model, which should lead to more accurate species delimitation using heuristic criteria such as *gdi.*

## HEURISTIC SPECIES DELIMITATION USING BPP

Here we describe how Bayesian parameter estimation under the MSC model can be combined with *gdi* to delimit species using a hierarchical procedure based on a species/population tree. This is similar to the use of a ‘guide tree’ for species delimitation by Yang and Rannala (2010), in that an ancestral node on the guide tree is merged into one species only if its descendant nodes are merged. However, here we rely on Bayesian parameter estimation on a fixed species/population tree while Yang and Rannala (2010) used reversible-jump algorithms to calculate posterior probabilities for different speciess delimitation models (represented by merging nodes on the guide tree). We first demonstrate the procedure using a simulated dataset and then apply it to the analysis of three empirical datasets analyzed previously by Jackson *et al.* (2017). The *gdi* is only one of many possible heuristics with rough correspondences to different species definitions.

We use a species/population tree for five populations, (*(((X,A),B),C),D*), to simulate data (Fig. 5a). *ABCD* represents a large paraphyletic species with a broad geographic distribution arranged in a stepping-stone design, with migration between any two adjacent populations including the ancestors (for example, between *D* and the ancestral population *XABC* after the first population split, and then between C and *D* and between C and *XAB* after the second split, etc.). The scaled migration rate is *M = Nm = 2* for any pair of adjacent populations. *X* is a new species, having separated from population *A* (with *τ_XA_* = 0.01), and there is no gene flow involving *X.* The divergence times (τs) are at 0.04, 0.03, 0.02, and 0.01. The population size parameter is *θ* = 0.01 for all populations. We simulated 100 loci, each of 500 sites, for four samples per species (20 sequences per locus).

To generate a working species/population tree (the guide tree), we run a joint analysis of species delimitation and species tree estimation (the A11 analysis in BPP, Yang, 2015). The parameters in the MSC model are assigned diffuse inverse-gamma priors *θ* ~ IG(3,0.02) and *τ* ~ IG(3,0.08), with shape parameter 3 and with the prior means matching the true values. We used a burnin of 40,000, sample frequency of 10, and collected 50,000 samples. We conducted four separate runs for each analysis, with convergence ensured mainly by checking consistency between runs. The posterior probabilities for the species delimitation models calculated in the A11 analysis provided strong support for five species, and the inferred species tree incorrectly placed species *X* sister to *ABCD* (Fig. 5b). This incorrect topology may be expected, as populations exchanging genes tend to form clades in species tree analyses that ignore migration (Leaché *et al.*, 2013). Next, we run an A00 analysis, estimating parameters on the inferred guide tree (Fig. 5b) to generate the posterior distribution for the *gdi* for the most recent species divergences, between *A* and *B* and between C and *D* (Fig. 5c). Note that *2τ_AB_*/ *θ_A_* is used to decide whether population *A* is a species distinct from *B*, while *2τ_AB_*/ *θ_B_* is used to decide whether population *B* is a species distinct from *A.* Low *gdi* values of < 0.2 indicate that *A* and *B* are one species, as are C and *D.* Next, we collapse *A* and *B*, and C and *D*, and conduct another A00 analysis to estimate *θ* and *τ* for putative species *AB* and *CD* (Fig. 5d). The posterior distribution of *gdi* obtained suggest that *AB* and *CD* belong to the same species (Fig. 5e). The final iteration fits a two-species model containing species *X* and species *ABCD* (Fig. 5I). The *gdi* value for species *ABCD* is ambiguous (with 0.2 < *gdi* < 0.7), while the evidence for species *X* is strong (*gdi >* 0.7, Fig. 5g). Here the *gdi* shows an ambiguity of the species status of *X* and *ABCD*, depending on which population size (*θ_X_* or *θ_ABCD_*) is used to calculate the index.

**FIGURE 5.**
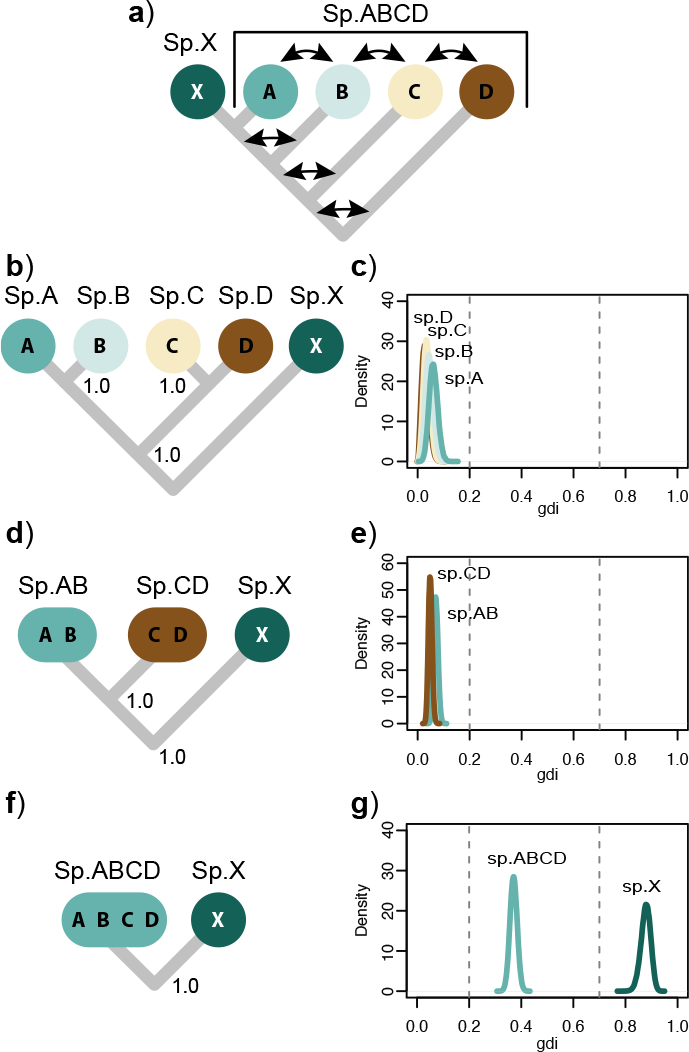
Species delimitation applying heuristic index *gdi* to parameter estimates from BPP. a) Species tree used for simulation allows migration between populations *A,B,C*, and *D* and their ancestors (indicated by arrows), but no gene flow involving species *X.* b) Species (guide) tree inferred from A11 analysis of BPP. In panels b–g, *gdi* is used to collapse populations on guide tree into same species in a hierarchical procedure, with BPP used to estimate MSC parameters (*θ* and *τ*) and generate posterior distribution of *gdi.* For example, *gdi* calculated using population *A* of panel b, based on *2τ_AB_/θ_A_* (equation 6), is shown in panel c (labeled ‘sp. *A’).* Sister populations inferred to belong to same species by *gdi* are collapsed, and resulting species tree is used to conduct a new BPP analysis. Procedure is repeated until distinct species are inferred or until root of tree is reached. According to Jackson *et al.* (2017), *gdi* < 0.2 indicates a single species, *gdi >* 0.7 indicates distinct species, and *gdi* values between 0.2 and 0.7 represent ambiguous species status.

Next, we re-analyzed the three empirical datasets of Jackson *et al.* (2017) using the hierarchical procedure described above. The three empirical datasets include eight nuclear loci from three populations of North American ground skinks (*Scincella lateralis*), 20 loci from three populations of southeastern United States pitcher plants (*Sarracenia alata*), and 50 loci from four population of *Homo sapiens.* In the analysis of Jackson *et al.* (2017), PHRAPL supported a single species of *Scincella lateralis* and two species of *Sarracenia alata*, and grouped the human populations into one species, while Bayesian model selection by BPP inferred the maximum number of species in each dataset.

Here we used the MCMC samples generated in the BPP analysis (Yang, 2015, analysis A00) to estimate the posterior distribution of the *gdi.* We used inverse-gamma priors on parameters (*θ*s and *τ*s), with the shape parameter 3 and with the same prior means as used by Jackson *et al.* (2017). For each dataset, we conducted four separate runs with a burnin of 10,000, sample frequency of 5, and collected 100,000 samples. The guide species trees are fixed at the previously published topologies from Jackson *et al.* (2017) (Fig. 6). We applied the hierarchical procedure to calculate *gdi* for population pairs by collapsing populations into a single species and conducting new MCMC analyses. Using BPP to calculate posterior distributions for *gdi*, we find no support for multiple species (*gdi >* 0.7) in any of the empirical datasets (Fig. 6).

## DISCUSSION

### Subjective allopatric species delimitation

We take it for granted that the neutral genome contains useful information about the population divergence history and about species status. In clear-cut cases, population divergence parameters should be sufficient to determine species status. For example, distantly related species can be reliably identified using a simple genetic distance threshold as in DNA-barcoding analysis (Hebert *et al.*, 2004). The difficulty is in identifying the species boundary (the so-called boundary conditions, Moritz and Cicero, 2004) for allopatric populations with low levels of genetic divergence and possibly frequent gene flow. The inherent subjectivity of allopatric species delimitation is clearly illustrated by the distinction between *statistical significance* and *biological significance* made by Jackson *et al.* (2017). Consider by analogy the estimation of the probability of heads in a coin-tossing experiment to determine the possible bias of the coin. Powerful statistical methods may detect a small bias in the coin, with *p* = 0.51, say. However, the bias of 0.01 is said to be statistically significant but not biologically significant, and it is considered incorrect to suggest that the coin with *p* = 0.51 is biased. Similarly, the definitions of races, subspecies and species are often subjective, and the neutral genome cannot provide unambiguous resolution of species status (Rannala, 2015).

### How to simulate if you must

The PSM specifies a process of population splits (incipient species formation) as well as conversions of such incipient species (populations) into true species. However, with time running forward, simulation under the PSM produces a new species (a conversion event) instantaneously. At a conversion event, the new true species and its parental incipient species (population) are deemed distinct species. As stated previously, this process does not realistically model the biological process, nor does it mimic the way taxonomists identify new species. We consider two alternative approaches for simulating the process of population splits and species assignments, and discuss their implications for the development of methods for species delimitation using genomic sequence data. A clear specification of the simulation procedure implies a probabilistic model of data generation and statistical inference methodology, because given the model, full-likelihood methods (maximum likelihood and Bayesian inference) are known to have certain statistical properties (Rannala, 2015).

**FIGURE 6.**
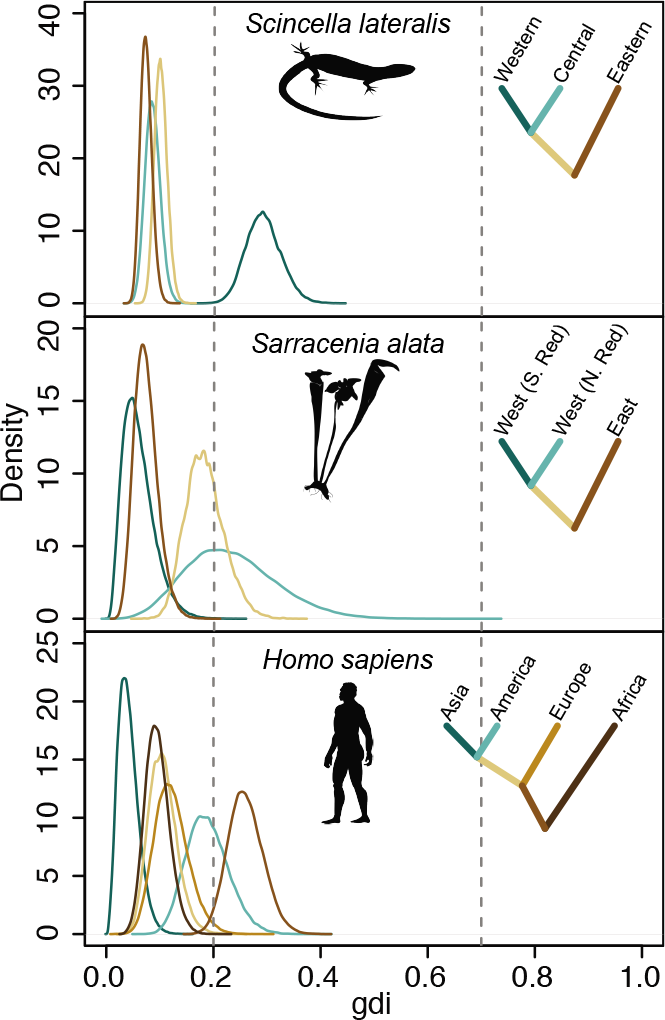
Posterior distribution of genealogical divergence index (*gdi*), generated in BPP analysis of three real datasets of Jackson *et al.* (2017). Silhouettes of species are from phylopic.org http://phylopic.org. Colored ancestral branches were analyzed by collapsing descendent species and conducting new MCMC analyses.

In the first approach, one can simulate population splits under a branching model, such as a variable-rate birth-death process. The random birth and extinction events specify a probabilistic distribution of the population tree topology and divergence times (*τ*s), and a certain model may be used to sample the population sizes (*θ*s) and migration rates (Ms). Gene trees (topologies and coalescent times) can then be generated using the population tree with parameters (*τ*s, *θ*s, Ms), and are then used to simulate sequence alignments. At the end of this simulation, the populations at the tips of the population phylogeny are assigned species status using heuristic criteria of divergence times and migration rates. This is very similar to the simulation approach of Jackson *et al.* (2017).

In the second approach, one may simulate population splits as in the first approach, but in addition simulate the evolution of a continuous character along the branches of the generated population phylogeny. The difference in the continuous character between two populations is a measure of genetic incompatibility and a threshold can be used to identify species status: if the continuous character has measurements *x_i_* and *x_j_* in two populations, they are considered distinct species if and only if |*x_i_* – *x_j_*| > *d*. The model for simulating the evolution of the continuous character may be a mixture of a small probability for ‘catastrophes’ (mimicking large events that may establish reproductive isolation at an instance, such as chromosomal rearrangements or polyploidizations) and a large probability for Brownian motion-like drift over time (mimicking the accumulation of genetic incompatibilities over time). At the end of the simulation, species status is assigned for populations at the tips of the tree based on the continuous character.

In both approaches, we assume that the process of sequence evolution is independent of population split events, and of the evolution of the continuous character, as expected if the neutral genome is used for species delimitation. Both scenarios seem to suggest that the only inference possible using the neutral genome is the population history and the population divergence parameters (*θ*s, *τ*s, and Ms). Assignment of species status will then depend on our empirical knowledge about the level of genetic divergence between good species, or the amount of genetic incompatibility that can be accumulated over a given time period.

### *The limited scope of* BPP *analyses*

The MSC model was developed for comparative analysis of the ‘neutral’ genome to estimate parameters that characterize the history of population divergences, under the assumption that natural selection has not significantly altered the genealogical histories of genomic regions (gene tree topologies and coalescent times). The MSC model does not aim to identify speciation genes or genes responsible for establishing reproductive barriers (which may be under species-specific directional selection), even though identifying such genes, however rare they are, may greatly enrich our understanding of the origin and maintenance of species. For example proteins involved in female and male reproduction are well-known to evolve at accelerated rates, apparently driven by natural selection due to ecological adaptations and sexual selection maintaining species boundaries (Swanson and Vacquier, 2002). In a few cases where the MSC model was applied to exons or the coding genome, it was noted to produce results highly consistent with the noncoding regions of the genome (Dalquen *et al.*, 2017; Ebersberger *et al.*, 2007; Shi and Yang, 2018). This is apparently due to the fact that most protein-coding genes are performing the same conserved functions in closely related species so that the effect of purifying selection removing nonsynonymous mutations is predominantly a reduction of the neutral mutation rate. The MSC model treats genomic regions as neutral markers to extract information concerning genealogical histories of the populations, reflected in population divergence parameters, such as population sizes, divergence times, and migration rates.

If species divergence is due to very few genes (in the so-called speciation islands) while the rest of the genome is homogenized due to widespread interbreeding, the overall divergence between species will be similar to the polymorphism within species (Nadeau *et al*., 2012). In such cases the neutral genome may not be informative about the species status and use of other kinds of data, such as evidence of ecological adaptation, or identification of speciation genes, etc., may be necessary to determine species status.

### Hypothesis tests versus parameter estimation

In this paper we have made a distinction between two kinds of analysis under the MSC model as implemented in BPP: (i) Bayesian model selection to calculate posterior probabilities for different species delimitation models (the A10 and A11 analyses in Yang, 2015) and (ii) Bayesian parameter estimation when species/population assignment and phylogeny are fixed (the A00 analysis in Yang, 2015). In theory, model selection can also be conducted in a Frequentist framework using a likelihood ratio test with the one-species model formulated to be the null hypothesis (with *τ* = 0) while the two-species model is the alternative hypothesis (with *τ* > 0). This is similar to testing the null hypothesis of a fair coin (with *p* = 1/2) against the alternative hypothesis of a biased coin (with *p* ≠ 1/2). With sufficient data, model selection can be very powerful in identifying population splits even if the age of the divergence event (*τ*) is very young. This is analogous to the use of a large number of coin tosses to detect a very small bias of heads versus tails.

We suggest that Bayesian model selection is appropriate for identifying morphologically cryptic species. Even if the genomic data or the BPP program cannot distinguish populations and species, the genetic distinctness of the populations signifies the presence of reproductive barriers or isolation mechanisms. There seems to be no controversy in assigning species status to populations that exist in sympatry and are genetically distinct.

Species delimitation by Bayesian parameter estimation aims to estimate the population-divergence parameters (θs, *τ*s, and Ms) and then apply a heuristic species definition, such as a minimum divergence criterion, 2*τ*/θ > 1, maximum migration M < 1, or the *gdi.* Using the coin-tossing analogy, this approach is like estimating the probability parameter *p* using the counts of heads and tails, and then applying whatever definition of bias one assumes heuristically.

The *gdi* attempts to use the overall genetic divergence between two populations affected by the combined effects of genetic isolation and gene flow. The index appears to have weaknesses. First, the criterion depends on the population divergence time relative to the population size (2*τ*/*θ_A_* in the case of no gene flow). If the population is established by a few founder individuals, *N_A_* and *θ_A_* may be very small, and the use of *gdi* may lead to claims of species status even if the populations diverged very recently. It may be necessary to consider the (absolute) population divergence (*τ*) (Yang and Rannala, 2014) as well as the divergence relative to the population size. Second, there may be ambiguity when the two populations concerned have drastically different sizes. If *N_A_* ≪ *N_B_*, the use of *gdi* may lead to the awkward solution that *A* is a distinct species from *B* (if one uses sequences *a*_1_,*a*_2_ and *b* to calculate the index) but *B* is not a distinct species from *A* (if one uses sequences *a*, *b*_1_ *b*_2_). This is the case in the analysis of the simulated data in Fig. 5g. Third, *gdi* has a large range of indecision (0.2-0.7), although this may reflect the arbitrary nature of species delimitation rather than a weakness of the index itself. We suggest there may be a need to propose heuristic criteria for species delimitation given the near absence of objective criteria.

### Concluding remarks

The MSC model and its implementation in BPP provides a powerful method for inferring population divergence histories and estimating evolutionary parameters using the fast-accumulating genomic sequence data. With accurate estimates of important population parameters, one can apply any empirical criterion for defining species that the evolutionary biologist entertains. For these reasons, the MSC model and BPP will continue to be essential tools in the analysis of genomic data to better understand biodiversity despite the fact that the interpretation of these results in assessing species status may be debated.

**TABLE 1.**
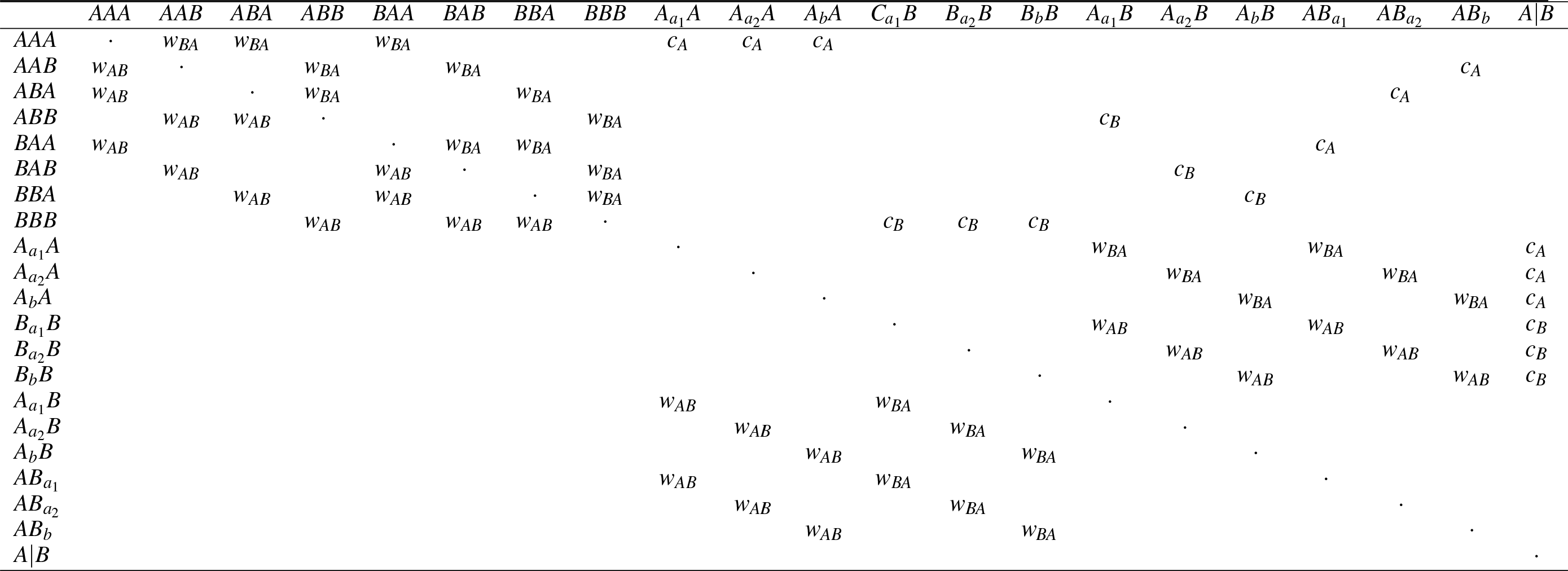
Rate matrix for Markov chain describing transitions between states in multispecies coalescent with migration model with two populations (*A* and *B*) and three sequences (*a*_1_, *a*_2_, and *b*).

## SUPPLEMENTARY MATERIALS

Data available from the Dryad Data Repository: http://dx.doi.org/???.

## ACKNOWLEDGMENTS

We thank N. Jackson for providing the R code for generating the PHRAPL results of Figure 4, and for clarifying the simulation design of Jackson *et al.* (2017). We thank F. Burbrink, B. Carstens, K. de Queiroz, N. Jackson, P. Kornilios, J. McGuire, and J. Mallet for their helpful comments. We are grateful to three anonymous reviewers for their constructive criticisms.

## REFERENCES

Burgess, R. and Yang, Z. 2008. Estimation of hominoid ancestral population sizes under Bayesian coalescent models incorporating mutation rate variation and sequencing errors. Mol. Biol. Evol., 25: 1979–1994.

Dalquen, D., Zhu, T., and Yang, Z. 2017. Maximum likelihood implementation of an isolation-with-migration model for three species. Syst. Biol., 66: 379–398.

Ebersberger, I., Galgoczy, P., Taudien, S., Taenzer, S., Platzer, M., and von Haeseler, A. 2007. Mapping human genetic ancestry. Molecular Biology and Evolution, 24: 2266–2276.

Edwards, S. V. 2009. Is a new and general theory of molecular systematics emerging? Evolution, 63: 119.

Etienne, R. S., Morlon, H., and Lambert, A. 2014. Estimating the duration of speciation from phylogenies. Evolution, 68(8): 2430–2440.

Hasegawa, M., Kishino, H., and Yano, T. 1985. Dating the human-ape splitting by a molecular clock of mitochondrial DNA. J. Mol. Evol., 22: 160–174.

Hebert, P. D., Stoeckle, M. Y., Zemlak, T. S., and Francis, C. M. 2004. Identification of birds through dna barcodes. PLoS Biol., 2: 16571663.

Heled, J. and Drummond, A. J. 2010. Bayesian inference of species trees from multilocus data. Mol. Biol. Evol., 27: 570–580.

Hey, J. 2010. Isolation with migration models for more than two populations. Mol. Biol. Evol., 27: 905–920.

Hobolth, A., Andersen, L., and Mailund, T. 2011. On computing the coalescence time density in an isolation-with-migration model with few samples. Genetics, 187: 1241–1243.

Jackson, N., Carstens, B., Morales, A., and B.C., O. 2017. Species delimitation with gene flow. Syst. Biol., 66: 799–812.

Jukes, T. and Cantor, C. 1969. Evolution of protein molecules. In H. Munro, editor, Mammalian Protein Metabolism, pages 21–123. Academic Press, New York.

Leaché, A. D., Harris, R. B., Rannala, B., and Yang, Z. 2013. The influence of gene flow on species tree estimation: a simulation study. Syst. Biol., 63(1): 17–30.

Maddison, W. 1997. Gene trees in species trees. Syst. Biol., 46: 523–536.

Mailund, T., Dutheil, J. Y., Hobolth, A., Lunter, G., and Schierup, M. H. 2012. Estimating divergence time and ancestral effective population size of Bornean and Sumatran orangutan subspecies using a coalescent hidden Markov model. PLoS Genet., 7(3): e1001319.

Mallet, J. 2013. Concepts of species. In S. Levin, editor, Encyclopedia of Biodiversity, volume 6, pages 679–691. Academic Press, Massachusetts.

Moritz, C. and Cicero, C. 2004. Dna barcoding: Promise and pitfalls. PLoS Biol., 2: e354.

Nadeau, N. J., Whibley, A., Jones, R. T., Davey, J. W., Dasmahapatra, K. K., Baxter, S. W., Quail, M. A., Joron, M., Ffrench-Constant, R. H., Blaxter, M. L., Mallet, J., and Jiggins, C. D. 2012. Genomic islands of divergence in hybridizing heliconius butterflies identified by large-scale targeted sequencing. Philos Trans. R. Soc. Lond. B. Biol. Sci., 367(1587): 343–353.

Nichols, R. 2001. Gene trees and species trees are not the same. Trends Ecol. Evol., 16: 358–364.

O’Hagan, A. and Forster, J. 2004. Kendall’s Advanced Theory of Statistics: Bayesian Inference. Arnold, London, England.

Pinho, C. and Hey, J. 2010. Divergence with gene flow: models and data. Ann. Rev. Ecol. Evol. Syst., 41: 215–230.

Rannala, B. and Yang, Z. 2003. Bayes estimation of species divergence times and ancestral population sizes using DNA sequences from multiple loci. Genetics, 164(4): 1645–1656.

Rannala, B. and Yang, Z. 2013. Improved reversible jump algorithms for Bayesian species delimitation. Genetics, 194: 245–253.

Rannala, B. H. 2015. The art and science of species delimitation. Curr. Zool., 61: 846–853.

Rosenblum, E., Sarver, B., Brown, J., Des Roches, S., Hardwick, K., Hether, T., Eastman, J., Pennell, M., and Harmon, L. 2012. Goldilocks meets Santa Rosalia: An ephemeral speciation model explains patterns of diversification across time scales. Evolut. Biol., 39: 255–261.

Shi, C. and Yang, Z. 2018. Coalescent-based analyses of genomic sequence data provide a robust resolution of phylogenetic relationships among major groups of gibbons. Molecular Biology and Evolution, 35: 159–179.

Sukumaran, J. and Knowles, L. 2017. Multispecies coalescent delimits structure, not species. Proc. Natl. Acad. Sci. USA., 114: 1607–1612.

Swanson, W. and Vacquier, V. 2002. The rapid evolution of reproductive proteins. Nature Rev. Genet., 3: 137–144.

Takahata, N., Satta, Y., and Klein, J. 1995. Divergence time and population size in the lineage leading to modern humans. Theor. Popul. Biol., 48: 198–221.

White, H. 1982. Maximum likelihood estimation of misspecified models. Econometrica, 50: 1–25.

Xu, B. and Yang, Z. 2016. Challenges in species tree estimation under the multispecies coalescent model. Genetics, 204: 1353–1368.

Yang, Z. 2002. Likelihood and Bayes estimation of ancestral population sizes in hominoids using data from multiple loci. Genetics, 162(4): 1811–1823.

Yang, Z. 2014. Molecular Evolution: A Statistical Approach. Oxford University Press, Oxford, England.

Yang, Z. 2015. The BPP program for species tree estimation and species delimitation. Curr. Zool., 61: 854–865.

Yang, Z. and Rannala, B. 2010. Bayesian species delimitation using multilocus sequence data. Proc. Natl. Acad. Sci. USA, 107: 9264–9269.

Yang, Z. and Rannala, B. 2014. Unguided species delimitation using DNA sequence data from multiple loci. Mol. Biol. Evol., 31: 3125–3135.

Yang, Z. and Rannala, B. 2017. Species identification by Bayesian fingerprinting: a powerful alternative to DNA barcoding. Mol. Ecol., 26: 3028–3036.

Yang, Z. and Zhu, T. 2018. Bayesian selection of misspecified models is overconifdent and causes spurious posterior probabilities for phylogenetic trees. Proc. Nat. Acad. Sci. USA.

Zhang, C., Zhang, D.-X., Zhu, T., and Yang, Z. 2011. Evaluation of a Bayesian coalescent method of species delimitation. Syst. Biol., 60: 747–761.

Zhu, T. and Yang, Z. 2012. Maximum likelihood implementation of an isolation-with-migration model with three species for testing speciation with gene flow. Mol. Biol. Evol., 29: 3131–3142.

